# Tree diversity increases productivity through enhancing structural complexity across mycorrhizal types

**DOI:** 10.1101/2023.04.11.536229

**Authors:** Tama Ray, Benjamin M. Delory, Helge Bruelheide, Nico Eisenhauer, Olga Ferlian, Julius Quosh, Goddert von Oheimb, Andreas Fichtner

## Abstract

Tree species diversity plays a central role for forest productivity, but factors driving positive biodiversity-productivity relationships remain poorly understood. In a biodiversity experiment manipulating tree diversity and mycorrhizal associations, we examined the roles of above- and belowground processes in modulating wood productivity in young temperate tree communities, as well as potential underlying mechanisms. We found that tree species richness increased forest productivity indirectly by enhancing structural complexity within communities. After six years, structurally complex communities were twice as productive as structurally simple stands. This pattern was consistent across stands with different mycorrhizal associations. Our results also demonstrate that taxonomic diversity and functional variation in shade tolerance, but not phenotypic plasticity, are key drivers of structural complexity in mixtures, which in turn lead to overyielding. Consideration of stand structural complexity appears to be a crucial element in predicting carbon sequestration in the early successional stages of mixed-species forests.

## Introduction

Numerous studies showed that forest productivity increases with tree species richness at local (*1*) and global scales (*2*). The mechanisms underlying these positive biodiversity-productivity relationships (BPRs) in forest ecosystems have been the focus of a range of tree biodiversity experiments set up in different biomes (*3*). Species interactions at the local neighbourhood scale, which can lead to reduced competition through niche partitioning or facilitation (*4*), spatial complementarity of tree crowns in canopy space due to neighbour-mediated shifts in crown traits and allocation patterns (*5, 6*), and temporal variation in functional traits within a community (*7*) have been identified as important drivers of overyielding in experimental tree communities (i.e., higher yield in a mixture compared to the weighted average of monoculture yields) (*8*). However, the role of stand structural complexity and functionally distinct mycorrhizal associations within a community in mediating BPRs remains unclear.

In forests, stand structural complexity is caused by a high variation in tree size and a high dissimilarity in the spatial arrangement of tree crowns. It has been quantified with several indices, often based on one-or two-dimensional stand structural attributes such as variation in tree diameter and tree height or stand density (*9, 10*). Exploring the role of forest structural complexity in regulating species interactions and stand productivity, however, requires an accurate quantification of the three-dimensional (3D) morphology of individual trees as well as the space occupied by growing trees (*11*). Novel approaches, such as terrestrial laser scanning (TLS), have been applied to characterise the structural complexity within a stand by summarising information extracted from point clouds into a stand structural complexity index (SSCI; (*12*)). This index captures different facets of the structural complexity of forest plots: (1) the vertical distribution of structural elements, such as stems, branches and leaves, and (2) the density of these elements in 3D space. SSCI therefore encapsulates information about the 3D heterogeneity in biomass distribution of all trees within a community (*13, 14*).

The structural complexity of a stand is shaped by tree species diversity (*15, 16*). Several studies using SSCI showed that stands become structurally more complex with increasing tree species richness (*14, 17, 18*) and that positive biodiversity effects on complexity become stronger over time (*14*). Stand structural complexity is strongly related to canopy space occupation. For example, differences in crown architecture among species and neighbourhood-driven plasticity within species (*19*), such as plastic changes in branch traits and branching patterns (*20*), contribute to greater canopy complexity in mixed species tree communities compared to monocultures. In particular, the physical complementarity between trees in the vertical and horizontal space of the canopy allows species-rich tree communities to exploit light resources more efficiently (*5, 21*), with shade tolerant and light-efficient species occupying lower canopy layers (*22*). This more efficient light-use in mixtures should lead to higher aboveground productivity in structurally more complex forest stands. This positive relationship between structural complexity and forest productivity has been demonstrated in natural forests using 2D structural complexity measures (*23, 24*) or canopy rugosity (*25*). However, we still have much to learn about how tree species richness modulates the relationship between stand structural complexity and forest productivity, especially when structural complexity is measured using state-of-the-art techniques that allow for detailed characterisation of tree biomass distribution in 3D space.

While these approaches allow us to assess the role of aboveground structural complexity for productivity, next to nothing is known about how belowground biotic interactions affect this relationship. Associations between mycorrhizal fungi and tree roots are an essential type of interaction in forests (*26–28*). Temperate tree species are either associated with ectomycorrhizal fungi (EM) belonging to Asco-or Basidiomycetes, arbuscular mycorrhizal (AM) fungi of the phylum Glomeromycota (*29*), or simultaneously with both EM and AM fungi (*30, 31*). While EM tree species generally benefit from greater nitrogen mobilisation from organic matter and enhanced organic and inorganic resource uptake (*32*), trees associated with AM fungi mainly benefit from a greater uptake of less mobile nutrients such as phosphorus (*26, 29*). Mycorrhizal associations with AM or EM fungi are known to influence tree productivity differently (*33–35*). For instance, Deng et al. (2023) found that differences in nutrient acquisition strategies affect the direction of BPRs, with positive effects of tree species richness on AM tree productivity, but negative effects for EM trees (*36*). Despite the importance of mycorrhizal associations in mediating tree productivity, the extent to which mycorrhizal associations influence the structural complexity of forest stands remains to be elucidated.

In this study, we took advantage of a tree diversity experiment called MyDiv. Established in 2015, the MyDiv experiment is located in eastern Germany and was designed to explore the role of mycorrhizal associations in shaping biodiversity-ecosystem functioning relationships in temperate tree communities (*37*). This unique experiment consists of two orthogonal gradients of tree species richness and mycorrhizal associations, where communities were planted with either AM tree species, EM tree species, or both along a gradient of tree species richness (monocultures, 2-species and 4-species mixtures). Here, we aimed at identifying the mechanisms that underlie BPRs in young temperate tree communities by disentangling the relative importance of stand structural complexity and mycorrhizal associations in modulating BPRs. First, we tested the hypothesis that the structural complexity and wood productivity of tree communities increase with tree species richness, and that mixtures composed of AM and EM tree species are most productive and structurally more complex. Second, we evaluated the extent to which positive BPRs are driven by stand structural complexity and tree mortality, and whether the proportion of AM and EM trees in a community affects wood productivity. We hypothesised that tree species richness indirectly affects community productivity by enhancing structural complexity and by decreasing tree mortality within a community. Furthermore, we expected that a lower proportion of AM and EM trees in a community, which occurs when switching from monocultures of AM or EM trees (100%) to mixed communities (50% AM and 50% EM trees), would have a positive effect on wood productivity. Third, we explored mechanistic links between tree species richness, structural complexity, overyielding, and functional characteristics of tree communities. We focused our analysis on four functional characteristics of tree communities: the community-weighted mean and functional dispersion of shade tolerance and plasticity in annual wood productivity. We hypothesised that the positive net effect of tree species richness on stand structural complexity is mediated by shade tolerance and phenotypic plasticity of tree species within a community. In particular, we expected biodiversity effects on stand structural complexity to be greater in species-rich tree communities composed of functionally dissimilar species. To test these hypotheses, we measured stand structural complexity (SSCI, sensu ref. (*12*)) in 2021 using terrestrial laser scanning, and quantified annual wood productivity (increment in stem volume from 2015 to 2021) by measuring the radial and longitudinal growth of individual trees. For each tree species included in our experiment, shade tolerance values were taken from the literature (*38*), and phenotypic plasticity was quantified as the extent to which wood productivity of each species was affected by tree species richness.

## Results

### Stand structural complexity and productivity increase with tree species richness, but are not affected by mycorrhizal associations

Stand structural complexity index (SSCI) increased with tree species richness (*P* = 0.002) and was on average 25% higher in 4-species mixtures compared to monocultures (Fig. 1A). The magnitude and direction of this relationship, however, were consistent across mycorrhizal associations (Table 1; Fig. 1B) and SSCI did not significantly vary among mycorrhizal associations either (Table 1; Fig. 1C). Across mixtures, the magnitude of tree species richness effects was strong (Hedges’ g: 0.95), and effect sizes were almost three times higher in 4-species than in 2-species mixtures (Hedges’ g: 1.49 and 0.57, respectively; Fig. S1).

**Fig. 1.**
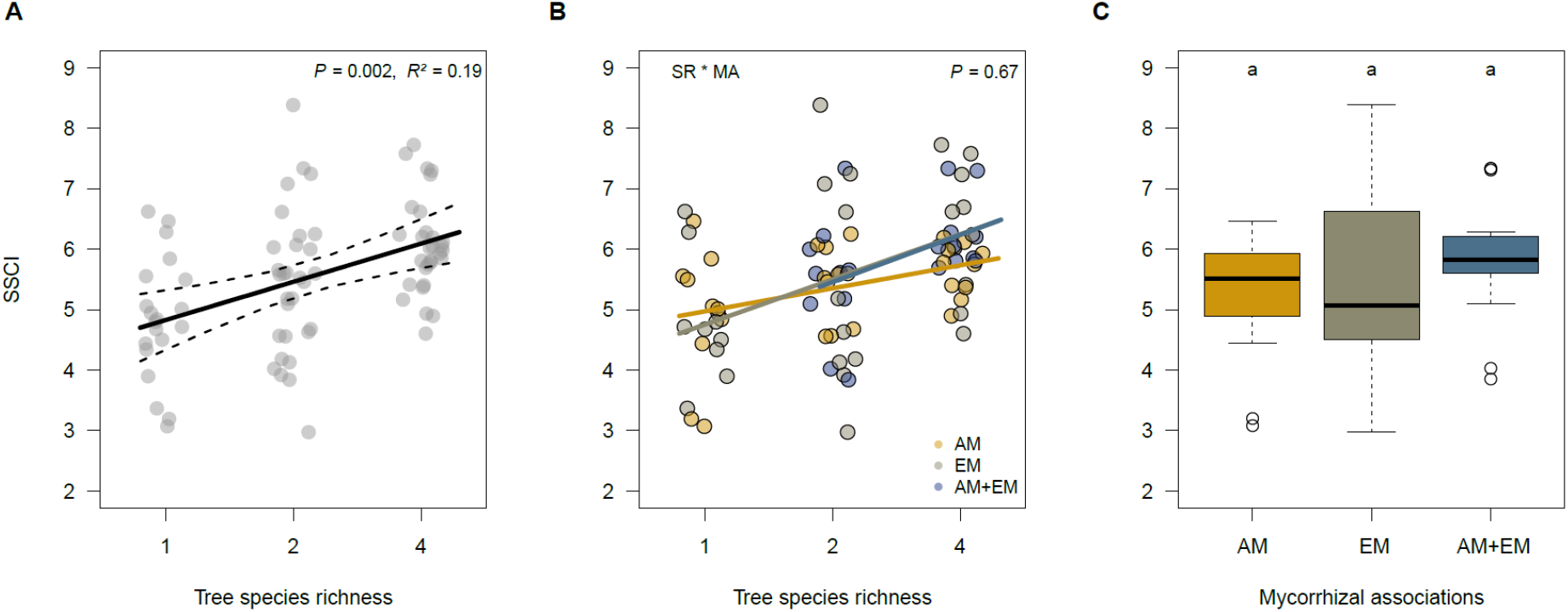
Tree diversity-stand complexity relationships. Panel A shows variations in stand structural complexity index (SSCI) as affected by tree species richness (SR) across mycorrhizal associations (MA), while panel B shows this relationship for each mycorrhizal association separately. The solid lines are mixed-effect model fits, and the dotted lines indicate the 95% confidence interval of the prediction. Individual dots represent the observed SSCI values, which are jittered to improve readability. The *R*²-value indicates the proportion of variance explained by SR alone. Panel C shows how SSCI was affected by mycorrhizal associations (across tree species richness levels). Boxplots show the median (horizontal black lines), the 25% and 75% percentiles (edges of the box) and 1.5 times the interquartile range (whiskers) of observed SSCI values. Open circles indicate SSCI values that are greater or smaller than 1.5 times the interquartile range. Differences among mycorrhizal associations were not statistically significant (Tukey-Test: *P* > 0.10).

**Table 1.**
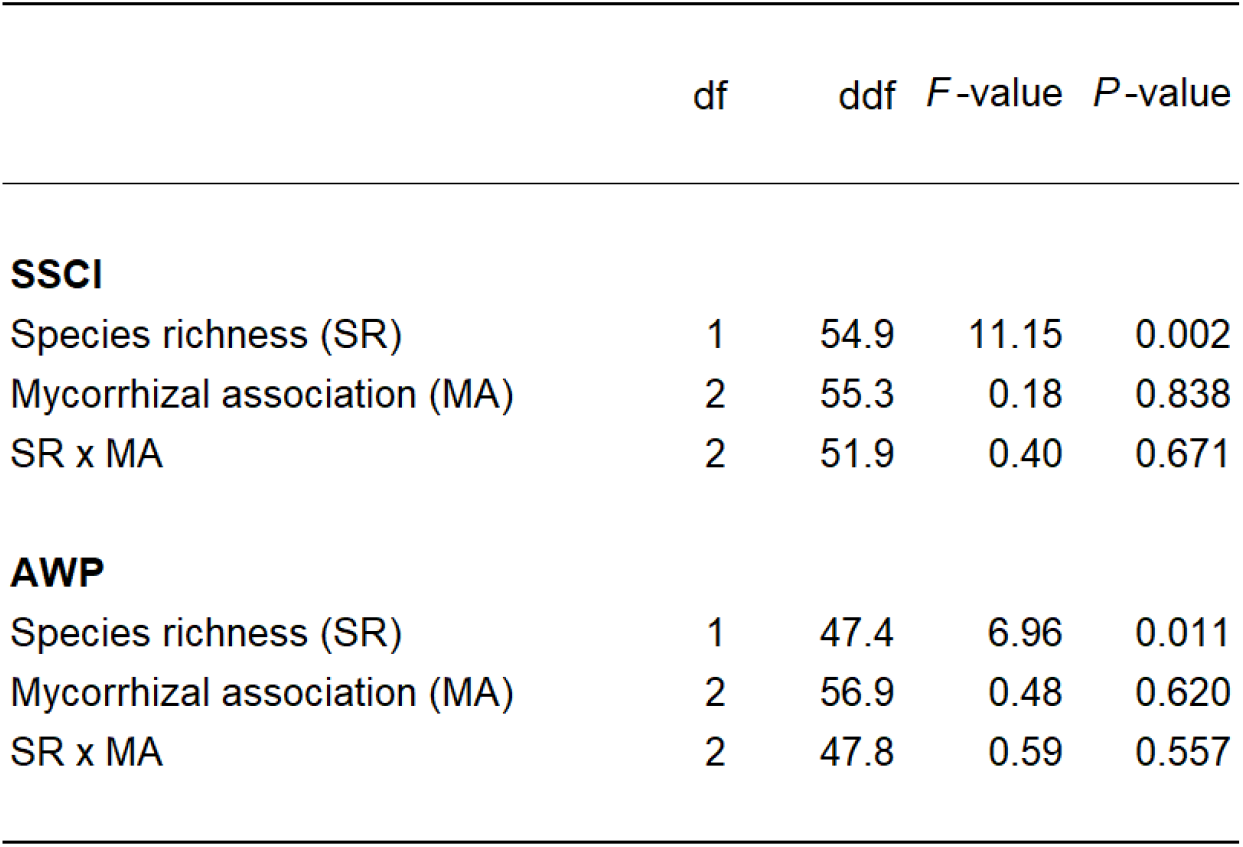
Effects of tree species richness and mycorrhizal associations on stand structural complexity (SSCI) and community productivity (AWP). Results of linear mixed-effects models using type-I sum of squares based on sequential fits of the fixed effects are indicated in the table. Tree species richness (log_2_SR) was fitted as a numeric variable, with mycorrhizal association being a categorical variable with three levels (AM, EM, AM+EM). SSCI, stand structural complexity index; AWP, annual wood productivity; df, numerator degrees of freedom; ddf, denominator degrees of freedom. See Table S1 for information on random effects and model fits of the best-fitted model.

|  | df | ddf | F-value | P-value |
| --- | --- | --- | --- | --- |
| **SSCI** |  |  |  |  |
| Species richness (SR) | 1 | 54.9 | 11.15 | 0.002 |
| Mycorrhizal association (MA) | 2 | 55.3 | 0.18 | 0.838 |
| SR x MA | 2 | 51.9 | 0.40 | 0.671 |
| **AWP** |  |  |  |  |
| Species richness (SR) | 1 | 47.4 | 6.96 | 0.011 |
| Mycorrhizal association (MA) | 2 | 56.9 | 0.48 | 0.620 |
| SR x MA | 2 | 47.8 | 0.59 | 0.557 |

Overall, the annual wood productivity (AWP) of tree communities increased with tree species richness (*P* = 0.01; Fig. 2A). After six years, 4-species mixtures accumulated an average of 28% more wood volume than monocultures. However, we found no support for the hypothesis that the relationship between tree species richness and AWP was modulated by mycorrhizal associations within tree communities (Table 1), despite the fact that communities with both arbuscular mycorrhizal (AM) and ectomycorrhizal (EM) tree species exhibited the strongest positive BPRs (Fig. 2B). Interestingly, AM communities shifted from being the most productive to being the least productive along the diversity gradient. In contrast, EM communities benefited most from growing in 4-species mixtures (Fig. S2), although these trends were not statistically significant (*P* > 0.05). Across tree communities, AWP did not differ significantly among mycorrhizal associations (all comparisons *P*_adj._ > 0.10), with variability in AWP being largest for EM communities (coefficient of variation: AM = 0.24, EM = 0.36; AM+EM: 0.25; Fig. 2C).

**Fig. 2.**
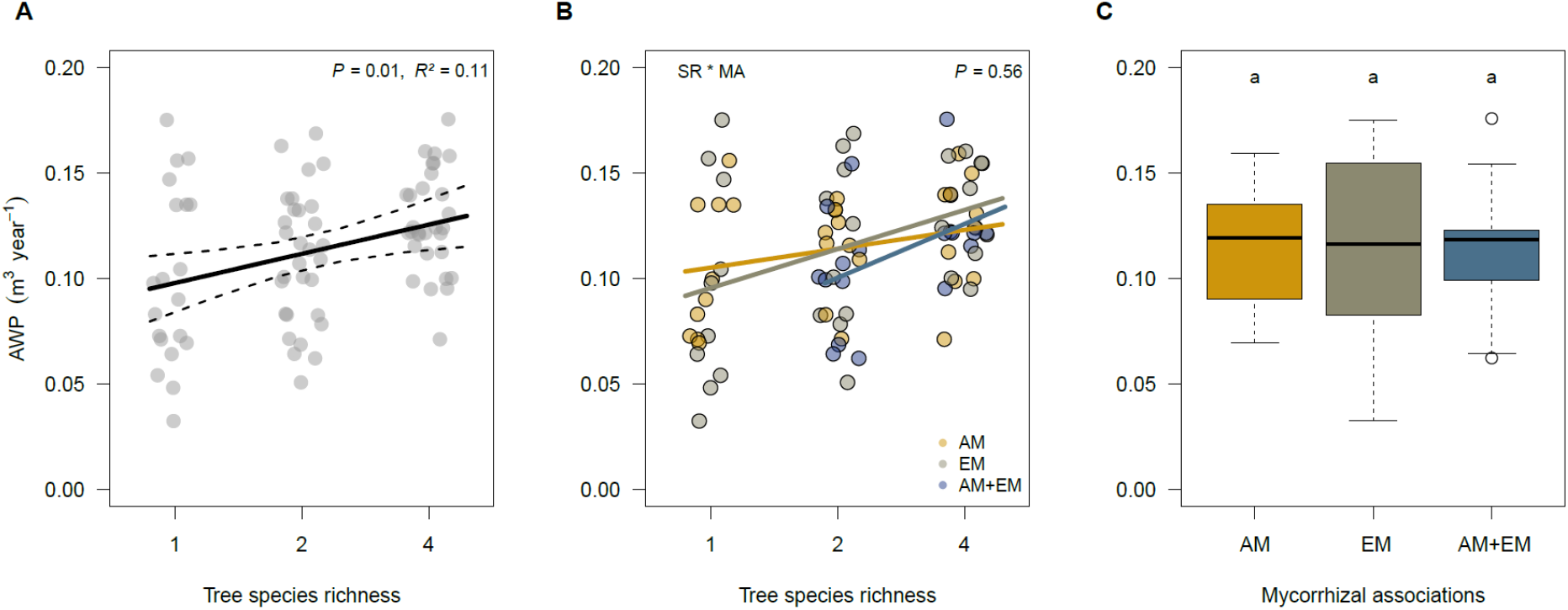
Tree diversity-productivity relationships. Panel A shows variations in annual wood productivity (AWP) as affected by tree species richness (SR) across mycorrhizal associations (MA), while panel B shows this relationship for each mycorrhizal association separately. The solid lines correspond to fitted biodiversity-productivity relationships of mixed-effects models, and the dashed lines indicate the 95% confidence interval of the predictions. Individual dots represent the observed AWP values, which are jittered to improve readability. The *R*²-value indicates the proportion of variance explained by SR alone. Panel C shows how AWP was affected by mycorrhizal associations (across tree species richness levels). Boxplots show the median (horizontal black lines), the 25% and 75% percentiles (edges of the box) and 1.5 times the interquartile range (whiskers) of observed community productivity. Open circles indicate productivity values that are greater or smaller than 1.5 times the interquartile range. Differences among mycorrhizal associations were not statistically significant (Tukey-Test: *P* > 0.10).

### Stand structural complexity modulates the relationship between biodiversity and productivity in young tree communities

Both monocultures and mixtures became more productive with increasing stand structural complexity (*P <* 0.001; Fig. 3). For example, the AWP of communities with a high structural complexity (95%-quantile of SSCI = 7.3) was almost two-fold (+97%) higher than those associated with a low structural complexity (5%-quantile of SSCI = 3.8).

**Fig. 3.**
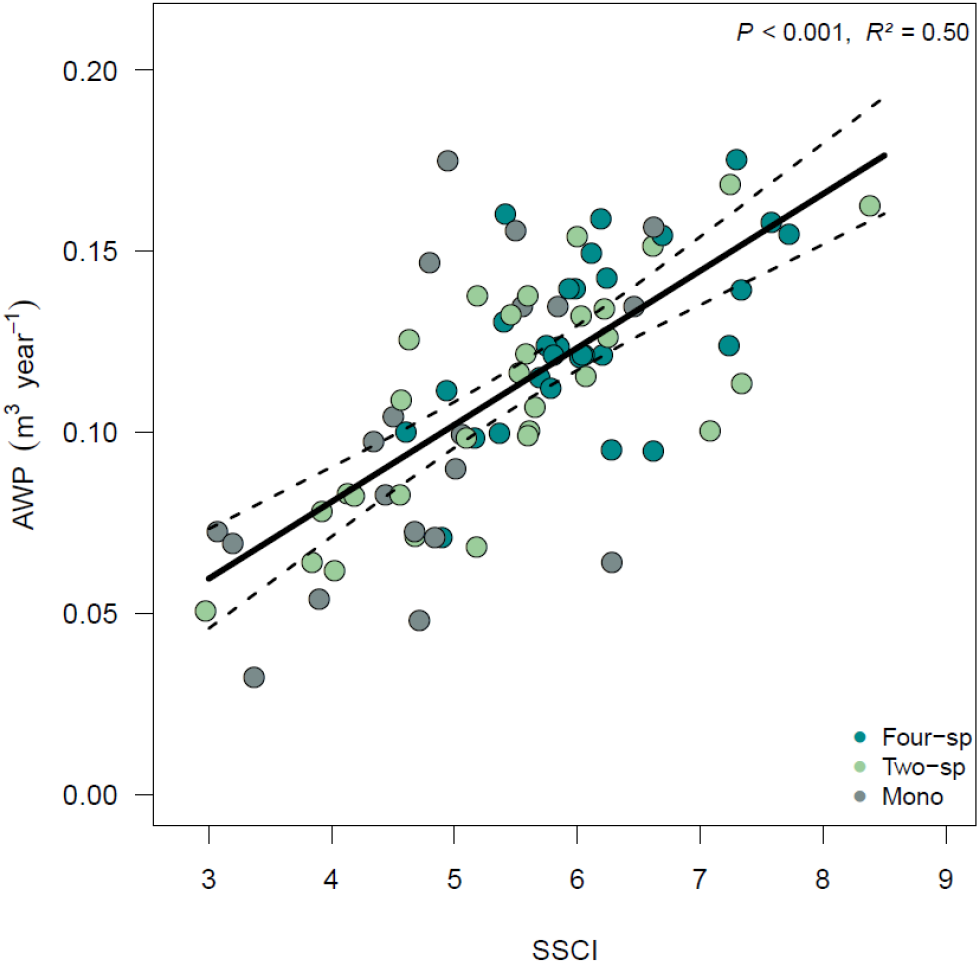
Relationship between stand structural complexity and wood productivity. Changes in annual wood productivity (AWP) with stand structural complexity index (SSCI) across tree communities. The solid line is a mixed-effect model fit, with dashed lines indicating the 95% confidence interval of the prediction. Dots represent AWP and SSCI values measured in each plot. *R*²-value refers to the proportion of variance in AWP explained by SSCI.

In order to understand how tree species richness and mycorrhizal associations affected tree mortality, structural complexity (SSCI), and wood productivity (AWP) in our young temperate forest plots, we fitted a first structural equation model (SEM) to our data (Fig. 4). SSCI, tree species richness, tree mortality, and mycorrhizal associations (i.e. proportion of AM and EM tree species in communities) explained 55% of the variation in AWP (Fig. 4). Importantly, we found that tree species richness indirectly increased AWP by enhancing structural complexity (*P* = 0.002), but not by mitigating tree mortality (*P* = 0.811). SSCI was the strongest driver of AWP. Even after controlling for other fixed effects in the productivity model and the influence of differences in tree species composition between communities, SSCI still explained a substantial amount (47.7%) of the variation in AWP. As expected, tree mortality had a negative effect on AWP (*P* = 0.008), but was not related to SSCI (*P* = 0.477). The relative importance of direct effects of tree mortality rate (*R*²: 5.0%), tree species richness (*R*²: 0.1%), and mycorrhizal associations (AM tree communities-*R*²: 1.2%; EM tree communities-*R*²: 0.1%) on AWP was small. We only found weak relationships between the proportion of AM and EM tree species in communities and AWP (AM: *P* = 0.258; EM: *P* = 0.738) (Fig. 4).

**Fig. 4.**
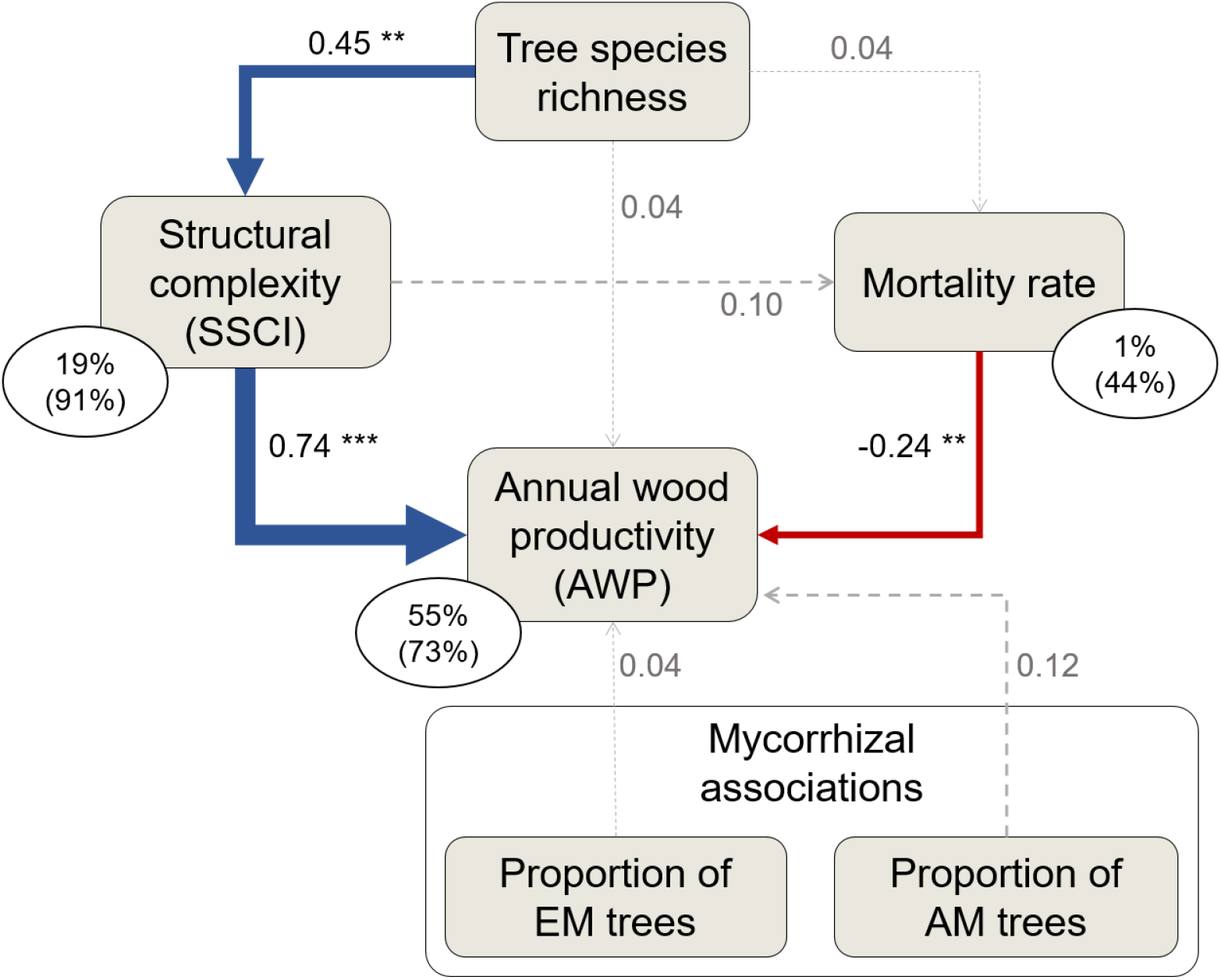
Structural equation model relating tree species richness, structural complexity, mortality rate, and mycorrhizal associations to annual wood productivity of tree communities. The blue and red arrows indicate significant (\**P* < 0.05, ** *P* < 0.01, *** *P* < 0.001) positive and negative relationships, respectively. Arrow width is proportional to standardised path coefficients. Dotted grey arrows indicate non-significant (*P* > 0.10) paths. Numbers next to arrows are standardised path coefficients. Percentage values are proportions of variance explained by fixed effects; proportions of variance explained by fixed and random effects are in parentheses. Note that communities with both arbuscular mycorrhizal (AM) and ectomycorrhizal (EM) tree species were coded as a reference in the SEM. This SEM fitted our data well (Fisher’s *C* = 9.41, *df* = 8, *P* = 0.309).

### Mechanisms underlying the positive relationship between tree species richness, structural complexity and wood productivity

For each mixture plot, we quantified the net effect of tree species diversity on (1) wood productivity (i.e., overyielding) and (2) stand structural complexity (ΔSSCI). The net biodiversity effect (NBE) on wood productivity was calculated and partitioned into complementarity and selection effects following (*8*). The net effect of tree species richness on stand structural complexity was estimated by calculating the difference between the SSCI measured by terrestrial laser scanning in mixed stands (SSCI_obs_) and the SSCI predicted based on the structural complexity measured in the monoculture plots of the constituent species (SSCI_pred_). A positive ΔSSCI value indicates that the structural complexity of a stand is greater than expected based on the complexity measured in the monoculture plots of its component species, while a negative value indicates the opposite. All tree mixtures had a greater productivity than expected based on monoculture yields (i.e., overyielding), and 87% of them were structurally more complex than would be expected based on the weighted average complexity measured in monoculture plots (i.e., positive values of ΔSSCI) (Fig. 6).

To understand the mechanisms underlying the positive biodiversity-complexity-productivity relationships in our experiment, we fitted two additional SEMs to our data. These SEMs separately investigated the relative importance of shade tolerance and phenotypic plasticity (community-weighted mean and functional dispersion values) in modulating the relationship between tree species richness and ΔSSCI (Fig. 5). For each species included in the MyDiv experiment, shade tolerance index (ST) values were taken from (*38*). We estimated phenotypic plasticity by quantifying the extent to which AWP of each species changed along the species richness gradient using the Relative Distance Plasticity Index (RDPI, (*39*)). Both SEMs fitted our data well (shade tolerance model in Fig. 5A: Fisher’s *C* = 3.08, *df* = 4, *P* = 0.545; phenotypic plasticity model in Fig. 5B: Fisher’s *C* = 1.88, *df* = 2, *P* = 0.391) and explained 36% and 43% of the variation in overyielding, respectively. We found that the net positive effect of tree species richness on structural complexity (ΔSSCI) was largely attributable to variation in taxonomic and functional attributes within communities, which in turn led to greater overyielding in mixtures (Fig. 5). This positive relationship between community overyielding and ΔSSCI was mostly driven by greater complementarity effects in structurally more complex tree communities (Fig. 6), suggesting an important role for species interactions in modulating complexity-productivity relationships in forests. In line with this, our results showed that interspecific variation in shade tolerance (FD_ST_) was a stronger determinant of ΔSSCI (*P* = 0.008, *R*²: 19%) than tree species richness (*P* = 0.012, *R*²: 11%; Fig. 5A). This indicates that species-rich communities composed of tree species with high and low shade tolerance were those associated with the strongest effects of biodiversity on structural complexity (Fig. S3), which then led to stronger biodiversity effects on wood production (Fig. 5A). In contrast, we found no evidence that variation in phenotypic plasticity between species (FD_RDPI_) modulated ΔSSCI in our experiment. However, we found that greater interspecific variation in tree plasticity had a direct negative effect on overyielding (*P* = 0.024; Fig. 5B). The community-weighted means of shade tolerance (CWM_ST_) and plasticity (CWM_RDPI_) had no significant direct effect on ΔSSCI (CWM_ST_: *P* = 0.270; CWM_RDPI_: *P* = 0.868). Tree species richness had no significant effect on the functional characteristics of communities, as neither the CWM nor FD values of shade tolerance and phenotypic plasticity were affected by tree species richness in our SEMs (Fig. 5).

**Fig. 5.**
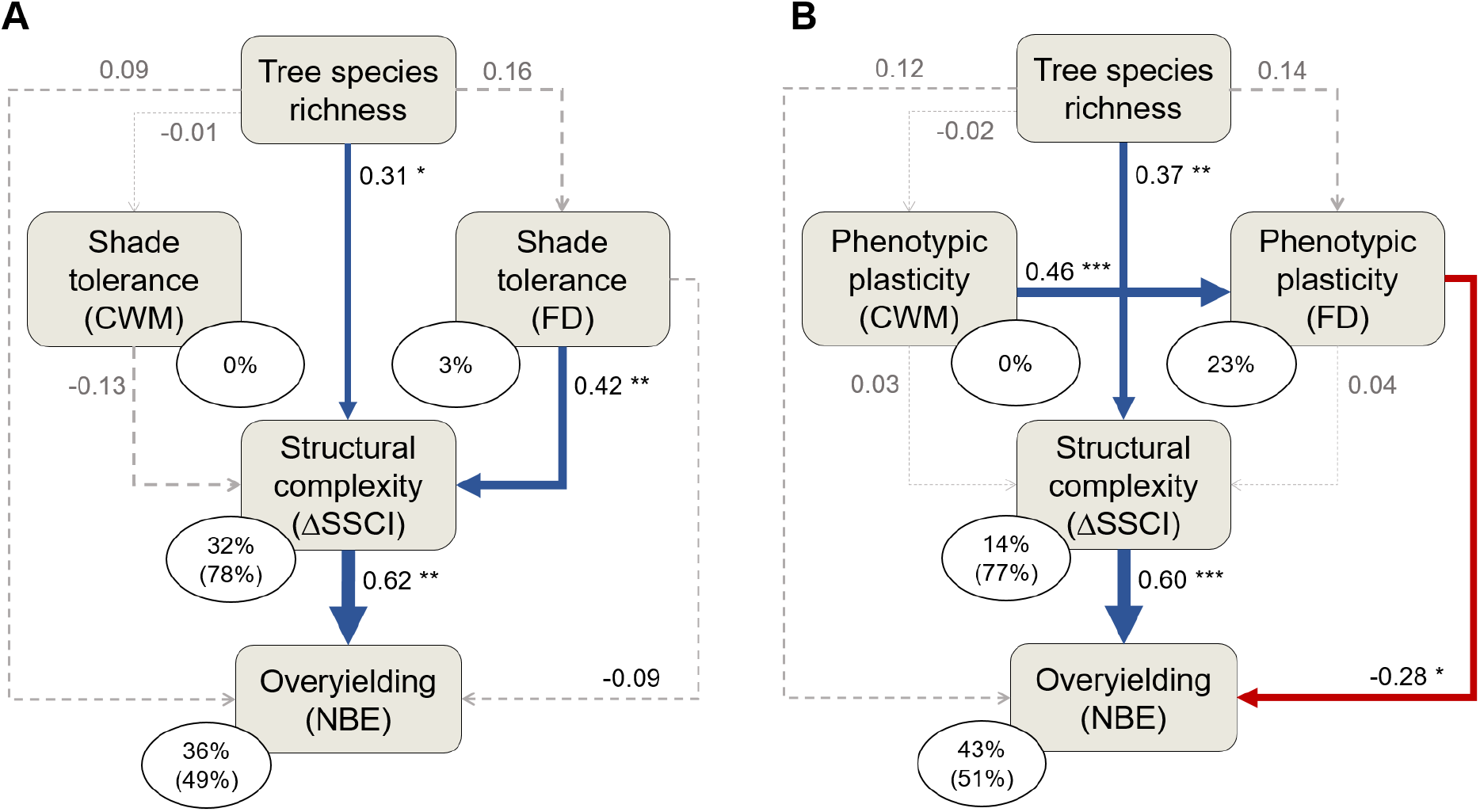
Structural equation models testing the roles of shade tolerance (A) and phenotypic plasticity (B) in mediating biodiversity-complexity-productivity relationships in young tree communities. For each mixed-species community, we calculated the community-weighted mean (CWM) and functional dispersion (FD) values of two key functional characteristics of tree species: (1) their shade tolerance (ST) and (2) their phenotypic plasticity in AWP (as measured by the Relative Distance Plasticity Index; RDPI). See Methods for more information about how these two functional characteristics of species were obtained or calculated. ΔSSCI represents the net effect of biodiversity on stand structural complexity (see Methods). The net biodiversity effect (NBE) on wood productivity was calculated following Loreau and Hector (2001) (*8*). The blue and red arrows indicate significant (\**P* < 0.05, ** *P* < 0.01, *** *P* < 0.001) positive and negative relationships, respectively. Arrow width is proportional to standardised path coefficients. Dotted grey arrows indicate non-significant (*P* > 0.10) paths. Numbers next to arrows are standardised path coefficients. Percentage values are proportions of variance explained by fixed effects; proportions of variance explained by fixed and random effects are in parentheses.

**Fig. 6.**
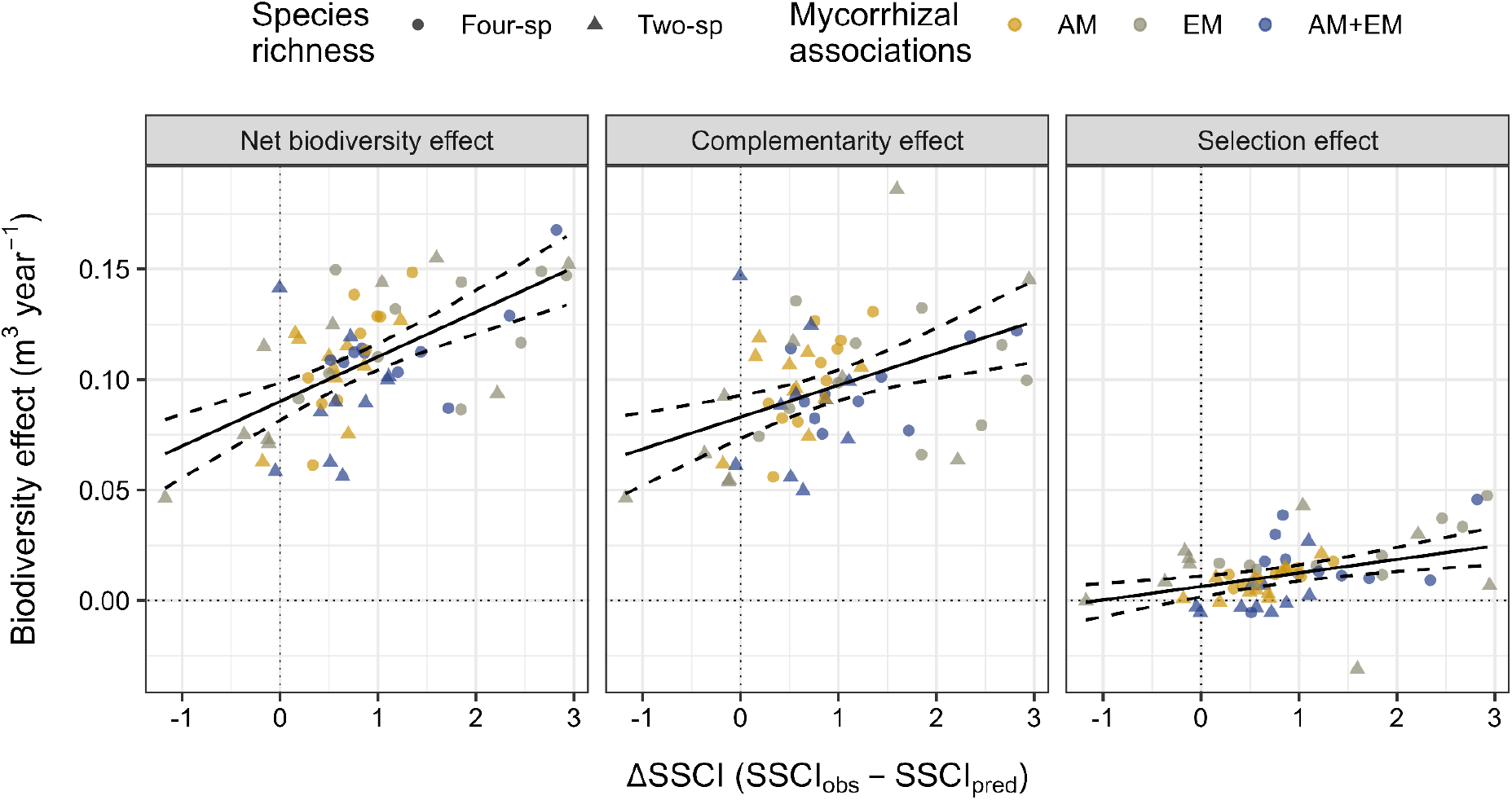
Overyielding in mixed-species communities is driven by stronger complementarity effects in structurally more complex tree communities. Dots are plot-level estimates of biodiversity effects on wood productivity and stand structural complexity (ΔSSCI). Solid black lines are predictions from mixed-effect models. Dashed lines indicate the 95% confidence interval of the prediction. ΔSSCI explains 35.6% (*P* < 0.0001), 18.0% (*P* = 0.0009) and 13.0% (*P* = 0.0016) of the variation in net biodiversity effects, complementarity effects, and selection effects, respectively.

The positive relationship between CWM_RDPI_ and FD_RDPI_ (Fig. 5B) was mainly brought about by the fact that EM tree species were more plastic than AM tree species (RDPI_EM_ > RDPI_AM_, see Table S2 and Fig. S4A). Because of this, AM and EM communities were characterised by low and high values of CWM_RDPI_ (Fig. S4B), respectively, but low FD_RDPI_ values compared to AM+EM mixtures (Fig. S4C). In contrast, mixtures of AM and EM tree species had intermediate CWM_RDPI_ values (Fig. S4B), but high FD_RDPI_ values (Fig. S4C).

Lastly, our results also support the existence of a productivity-plasticity trade-off, since we found that fast-growing species had a lower plasticity in AWP than slow-growing species (*P* = 0.049, *R*² = 0.40, Table S2, Fig. S5).

## Discussion

The MyDiv tree biodiversity experiment allowed us to quantify the relative importance of aboveground and belowground processes in driving biodiversity-productivity relationships (BPRs) in young temperate forests. Our results show that stand structural complexity is a fundamental driver of positive BPRs in forests with different mycorrhizal associations. The positive and strong biodiversity-complexity-productivity relationship in mixed-species tree communities was primarily due to increased taxonomic diversity and complementary light-capture strategies among shade-tolerant and shade-intolerant species, but not to differences in phenotypic plasticity among tree species.

As hypothesised, stand structural complexity and annual wood productivity increased with tree species richness. However, we found no evidence to support our hypothesis that the strength of BPRs depends on mycorrhizal associations, indicating that mixing trees with different types of mycorrhizae does not necessarily lead to a greater increase in wood productivity in diverse communities, at least in the first six years after establishment. Previous studies in mature forests and long-term tree diversity experiments have shown that the positive effects of tree species richness on forest productivity depend on the proportion of different mycorrhizal associations within a community (*35, 36*). However, the young developmental stage of our tree communities, nutrient legacies from previous agricultural use, and temporal variation in the importance of AM and EM trees in contributing to productivity in young mixed species communities (*33*) may explain why the effects of tree diversity on structural complexity and productivity were consistent across ecological strategies in our study.

We hypothesised that tree species richness had an indirect positive effect on community productivity by (1) mitigating tree mortality and (2) enhancing stand structural complexity. Although we found that plots with lower mortality rates were on average more productive, we found no relationship between tree species richness and tree mortality, suggesting that the expected reduction in tree mortality as a result of reduced intraspecific competition in mixtures did not occur during the first six years of our experiment (*40*). Given that plot-level mortality was comparatively low on average (median: 3.9%; mean: 7.7%), this may explain why we could not observe any significant mitigating effects of tree species richness on tree mortality. Instead, our results strongly support the hypothesis that community productivity is related to diversity-mediated shifts in stand structural complexity. We showed that the positive effects of tree diversity on wood productivity was strongly mediated by structural complexity within a stand: the higher the number of tree species in a plot, the greater the structural complexity and wood productivity of a forest stand. Although previous studies from tropical (*18*), subtropical (*14*), and temperate forests (*17*) using SSCI values derived from terrestrial laser scanning data have also demonstrated that tree species richness promotes structurally complex stands, our findings provide experimental evidence that variation in stand structural complexity acts as a central mechanism shaping BPRs in young tree communities. The greater structural complexity in mixtures also comes from the phenotypic plasticity of trees, as trees growing in species-rich neighbourhoods can adapt their crown morphology, both in size and shape, to maximise light capture, resulting in greater crown complementarity and wood volume growth (*5*). Therefore, community-level productivity should increase, as constituent species share canopy space more efficiently at the local neighbourhood scale, which may explain why community productivity was strongly and positively related to stand structural complexity in our study. Although niche partitioning in mixtures generally reduces the intensity of competition among individuals, we did not observe a reduction in tree mortality rates in forest stands with a greater structural complexity.

Our results showed that tree species richness and functional diversity in shade tolerance did not directly increase overyielding, but indirectly via enhancing the structural complexity of tree communities. In contrast, we found that a higher variation in phenotypic plasticity among tree species had a direct negative effect on overyielding. The strong positive relationship between tree species richness, stand structural complexity and wood productivity suggests that variation in phenotypic plasticity among species play a minor role in regulating productivity in early-successional stages. This is most likely the result of the trade-off observed between plasticity and productivity in our study, as fast-growing species in monocultures were associated with lower plasticity on average and vice versa. The existence of this trade-off means that mixing tree species with high and low phenotypic plasticity also means mixing tree species with low and high productivity, respectively, which does result in lower wood production than those of stands dominated only by productive (and, thus, less plastic) species. Furthermore, given that EM tree species were twice as plastic but 28% less productive than AM tree species, it is conceivable that species’ plastic responses to changes in tree species richness (*5, 41*) could have been affected by mycorrhizal associations.

Our finding that wood volume overyielding (i.e. net biodiversity effect on annual wood productivity) was strongly controlled by stand structural complexity can be largely attributed to net biodiversity effects on structural complexity (ΔSSCI). In our experiment, 52 out of 60 mixture plots were structurally more complex than expected based on the weighted average complexity of the monoculture plots of their constituent species. Given that high CWM values of phenotypic plasticity or shade tolerance (i.e. communities dominated by highly plastic or shade tolerant species) did not translate into stronger biodiversity effects on complexity, directional shifts in the functional composition of tree communities did not explain the observed increase in stand structural complexity with increasing tree species richness. This contradicts predictions from the mass-ratio theory (*42*), but suggests that mixing functionally different species is critical for shaping structural complexity in mixtures. In support of this, our results also highlight that mixing a greater number of tree species with different shade tolerance (i.e., increasing the taxonomic and functional diversity of a stand) enhances the net effect of biodiversity on complexity, with the increase in functional diversity being the most important in driving this effect. This is in agreement with several studies that have reported that dissimilarity in shade tolerance can increase forest productivity (*43–46*), suggesting that positive BPRs can result from greater complementarity in light use strategies (*47*).

Shade tolerance is a key ecological trait that shapes plant-plant interactions, because it is closely associated with functional traits related to a tree’s competitive ability for light (*48, 49*). For instance, fast-growing species are often shade-intolerant and vice versa (*50*), allowing shade-intolerant species to compensate for their lower ability to sustain competition by reaching the upper canopy layers more quickly (*51*). A higher variation in shade tolerance among species within a forest stand likely leads to more vertically structured canopies (*43*), where shade-intolerant species dominate the upper canopy layers, while shade-tolerant species populate lower canopy layers (*52*). This is in agreement with our finding that, among the tree species included in our experiment, those with greater shade tolerance were on average shorter than shade-intolerant species (Fig. S7). This relationship was primarily driven by two ectomycorrhizal species with contrasting life history strategies: *Betula pendula*, fast-growing and shade-intolerant, and *Fagus sylvatica*, slow-growing and shade-tolerant ((*33*); Fig. S7). These differences between *B. pendula* and *F. sylvatica* caused greater vertical stratification in stands where both species were present, which led to stronger biodiversity effects on structural complexity (Fig. S3) and wood productivity.

Importantly, our results showed that overyielding in mixed-species communities is primarily driven by stronger complementarity effects in structurally more complex tree communities, with selection effects playing a minor role. Greater complementarity in structurally more complex forest stands could be the result of more effective partitioning of light resources in patches with higher tree diversity. As a result, tree species growing in species-rich communities can occupy different sections along a light availability gradient, thereby increasing light interception and light use efficiency of tree communities, which in turn increases productivity (*22*). Our results suggest that complexity-dependent interactions between tree species for light exploitation play an important role for complementarity effects to arise. In addition to species dissimilarity in shade tolerance, other components of functional diversity related to light capture, such as interspecific differences in tree branching and architecture, crown traits, or leaf phenology, may also have contributed to the net effect of biodiversity on stand structural complexity (*15, 20, 53*), which could also explain the importance of taxonomic diversity in mediating structural complexity in the forest plots of the MyDiv experiment.

Our results highlight the importance of selecting species with contrasting functional traits associated with light capture rather than contrasting mycorrhizal types in forest restoration projects to increase biomass accumulation during early stand development. Structurally complex stands can also support greater biodiversity, as the amount of niche opportunities for other trophic levels increases with heterogeneity in the horizontal and vertical structure of the forest (*54, 55*). Therefore, improving stand structural complexity would benefit both biodiversity conservation and climate change mitigation, which is particularly relevant in the context of the UN Decade of (Forest) Restoration.

## Materials and Methods

### Study site and experimental design

The study was conducted within the MyDiv biodiversity-ecosystem functioning experiment (*37*). The experimental site is located in Bad Lauchstädt (51°230N, 11°530E, Germany) at 114–116 m a.s.l with a continental summer-dry climate. The mean annual temperature and the mean annual precipitation are 8.8°C and 484 mm, respectively. Soils are developed on a parent material of silt over calcareous silt (loess) and the soil type is a Haplic Chernozem with a thick humus horizon. The study site is a former agricultural land. It was used as arable land for centuries until 2012 and then converted to grassland from 2013 to 2015. Due to these agricultural legacies and pedogenesis, the soil at our experimental site has a high nutrient content, particularly in nitrogen (*37*).

The MyDiv experiment was set up using a species pool of 10 native deciduous temperate tree species. These species were selected to represent two main types of mycorrhizal associations, with five arbuscular mycorrhizal (AM) tree species and five ectomycorrhizal (EM) tree species (Table S2). Two main factors have been factorially manipulated in this experiment: tree species richness (three levels: 1, 2 or 4 species per plot) and mycorrhizal associations (three levels: only AM tree species, only EM tree species, or mixture of AM and EM tree species). The experiment consists of 80 plots of 121 m² (11 × 11 m) arranged in two blocks. In March 2015, each plot was planted with 140 trees using a regular planting distance of 1 m. At the time of planting, all tree individuals were two to three years old. The tree diversity gradient included monocultures (n = 20), 2-(n = 30), and 4-species mixtures (n = 30). For each species, monocultures were replicated twice. The species composition of 2-species mixtures was not replicated (i.e., all 2-species mixture plots had a unique species composition). In 4-species mixtures with only one mycorrhizal type (AM or EM), all possible species combinations have been implemented twice. In 4-species mixtures with both mycorrhizal types (AM+EM), however, only 10 unique species combinations were implemented. This experimental design resulted in 30 plots with AM tree species (n = 10 for each diversity level), 30 plots with EM tree species (n = 10 for each diversity level) and 20 plots with both mycorrhizal types (each n = 10 for 2-and 4-species mixtures). To ensure that all species were equally represented within a plot, individuals were planted in the same proportions and at the same overall density, with 70 (2-species mixtures) and 35 (4-species mixtures) individuals per species per plot. More details about the experimental design can be found in (*37*).

### Terrestrial laser scanning (TLS)

TLS data was collected with a Riegel VZ400i terrestrial laser scanner (Riegl, Horn, Austria) in September 2021 under leaf-on conditions.

The scanner was set up on a tripod at a height of 1.3 m and positioned at the centre of each plot. Two scans were taken per position to get a full spherical view. After taking the first vertical scan, the scanner was tilted by 90 degrees and the horizontal scan was performed. The angular resolution of the scanner was set to 0.04, which corresponds to a resolution of 7 mm at a 10 m distance with a laser frequency of 600 kHz. We chose these settings following Reich et al. (2021) to have a better canopy penetration. All scans were taken under a clear sky in almost windless conditions.

The point clouds of the two scans of each centre position were registered using Riscan pro 10.11.3 (http://www.riegl.com) and then clipped to one specific plot to remove information from the other plots. Filtering was conducted before registration and stray and noise points were removed based on pulse shape deviation and relative reflectance. In our study, all points with pulse shape deviation above 15 or reflectance less than 15 dB were removed following manufacturer’s instructions to improve the quality of the point clouds (*56*). Because many trees already branched near the soil surface, we chose not to remove all points below 1.3 m (as done by (*12*)), but to remove only the soil surface layer of each plot.

### Quantifying the structural complexity of tree communities

For each plot, an index of stand structural complexity (SSCI) was calculated based on a point cloud using the approach of (*12*). This approach has been found to provide an effective measure to quantify the structural heterogeneity and complexity of a forest stand (*13, 57*). The point cloud was converted to a voxel grid and then the ratio of the total number of filled voxels to the total number of voxels was quantified (*58*). We used a voxel size of 5 cm and a slice thickness of 25 cm following (*14*). SSCI consists of two components (Equation 1, Fig. S8): the effective number of layers (ENL) and the mean value of the fractal dimension of several cross-sections of the point cloud (MeanFrac).

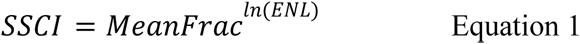

MeanFrac is a fractal dimension index of the cross-sectional polygon, derived from a 3D point cloud. It is dependent on the density of structural elements such as tree branches (*12, 13*). MeanFrac was calculated as the mean value of the fractal dimension of 4500 cross-sections. All points of each cross-section were combined in a polygon after sorting by angle. ENL quantifies the effective number of layers within a cross-section. It is closely related to the height of a tree and increases with increasing stand height (*14, 57*). Thus, it captures the vertical stratification within a community. ENL was calculated using the inverse Simpson-Index. We computed MeanFrac, ENL and SSCI using R 4.2.1 (*59*) with the packages VoxR (*60*) and sp (*61*).

### Tree growth data, mortality rate and biodiversity effects

To avoid edge effects, growth analyses were focused on the 64 trees growing in the centre (8 × 8 m) of each of the 80 study plots. For each tree *i*, we measured stem diameter (*D_i_*, measured in metres at 5 cm aboveground) and tree height (*H_i_*, measured in metres as the distance between the stem base and the apical meristem). For each tree, wood volume (*V_i_*, m³) was then estimated as 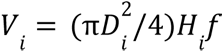 (Equation 2), where *f* is a cylindrical form factor of 0.5 (an average value for young broadleaved trees) to account for the deviation of the tree volume from the theoretical volume of a cylinder. For each plot *j*, annual wood productivity (AWP, m³ year^-1^) was calculated as the sum of the annual growth rates of all living trees (*n*) within a plot (Equation 3). In Equation 3, *V_ij,1_* and *V_ij,2_* are the wood volumes of tree *i* in plot *j* at the beginning (t_1_) and at the end (t_2_) of the growing period (2015–2021).

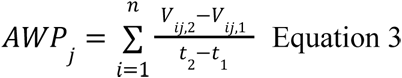

Plot-level mortality was quantified as the relative mortality of all trees within a plot between 2015 and 2021.

The net effect of biodiversity on AWP (NBE, also referred to as overyielding), as well as complementarity and selection effects, were calculated using the additive partitioning method of Loreau and Hector (2001) (*8*) implemented in the *bef* R package (*62*) available from GitHub (https://github.com/BenjaminDelory/bef).

### Quantifying the net effect of tree species diversity on stand structural complexity

The net effect of tree species diversity on stand structural complexity in plot *j* (*ΔSSCI_j_*) was calculated using Equation 4, where SSCI*_j,obs_* is the SSCI measured in plot *j* by TLS, SSCI*_j,pred_*is the SSCI value of plot *j* predicted based on the structural complexity measured in the monoculture plots of the species present in plot *j*, *S* is the tree species richness in plot *j*, *p_ij_* is the relative abundance of species *i* in plot *j* in 2021 (taking into account tree mortality from 2015 to 2021), and 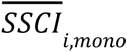 is the average SSCI measured in monocultures of species *i*. Positive ΔSSCI values indicate that the structural complexity of a stand is greater than expected based on the complexity measured in the monoculture plots of its component species, while negative values indicate the opposite. Typically, positive ΔSSCI values are expected if tree species richness has a positive effect on stand structural complexity.

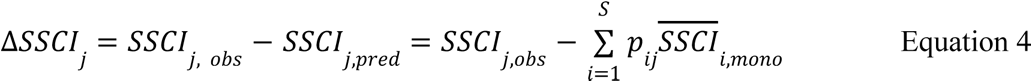

### Quantifying tree phenotypic plasticity

We assessed tree phenotypic plasticity by quantifying the extent to which AWP of each species changes along the species richness gradient (Fig. S5). We expect tree species with low phenotypic plasticity to show small differences in AWP between monoculture and mixtures of two and four species. Tree species with high phenotypic plasticity, however, would show large differences in AWP across species richness levels. Phenotypic plasticity was assessed using the Relative Distance Plasticity Index (RDPI) described in (*39*). For each tree species, RDPI was calculated as the mean relative difference in AWP between two plots (*j* and *j’*) with different richness levels (*i* and *i’*, with *i*≠*i*’) (Equation 5). In equation 5, *AWP_ij_*is the average tree productivity of a species in plot *j* (with species richness level *i*), and *N* is the total number of relative distances to calculate. *N* is calculated using Equation 6, where *n_1_*, *n_2_* and *n_4_* are the total number of plots in which a species grows in monoculture or with one or three other species, respectively. RDPI is a metric bounded between 0 (no plasticity) and 1 (maximal plasticity).

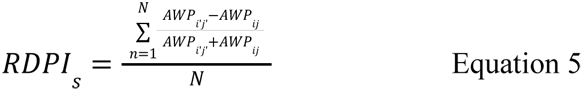

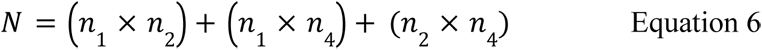

### Functional characteristics of tree communities

To understand better how tree species richness affects stand structural complexity (as measured by SSCI), we gathered data on two important functional characteristics for each tree species in our experiment: (1) their capacity to tolerate shade (as measured by a shade tolerance index, ST) and (2) their phenotypic plasticity in AWP (as measured by RDPI). ST values were taken from (*38*), while RDPI values were calculated as described above (Table S2). Species with high shade tolerance and phenotypic plasticity should have higher values for both indices.

For each tree community, we calculated the community-weighted mean and functional dispersion for shade tolerance (CWM_ST_ and FD_ST_) and phenotypic plasticity (CWM_RDPI_ and FD_RDPI_). These functional diversity metrics were calculated using the *FD* R package (*63*). Planting densities corrected for tree mortality between 2015 and 2021 were used to compute CWM and FD values.

### Statistical analysis

Prior to model fitting, tree species richness (SR) was log_2_-transformed and communities’ tree mortality rates (%) measured between 2015 and 2021 was square root-transformed to meet model assumptions, which were visually checked and confirmed according to (*64*). We used linear mixed-effects models to assess the impact of SR (numeric variable with 3 levels), mycorrhizal associations (MA; categorical, 3 levels: AM, EM, AM+EM) and their interacting effect on stand structural complexity (SSCI). Tree species composition of the plots was used as a random effect to avoid confounding effects between SR and compositional differences in tree species among communities. Preliminary analyses indicated that there was no significant difference in SSCI between the two experimental blocks (type-I sum of squares: *F* = 2.66, *P* = 0.117). Therefore, we did not use the study block as an additional covariate in the models. We used Hedges’ g effect size (*65*) as a standardized measure to quantify the strength of biodiversity effects (i.e., SR effects) on SSCI. Effect sizes were calculated based on predicted values of the best-fitted model at each level of tree species richness (monocultures, 2-and 4-species mixtures) using pooled standard deviation. Positive values of Hedges’ g indicate positive biodiversity effects on SSCI, and vice versa. Small, moderate and large effects are indicated by Hedges’ g values of 0.2, 0.5 and 0.8 (*66*).

To test the hypothesis that biodiversity-productivity relationships are modulated by mycorrhizal associations, we modelled AWP as a function of SR and MA, and their interaction as fixed effects. This linear mixed-effect model was fitted using tree species composition of the plots as a random effect. All linear mixed-effects models were fitted with restricted maximum likelihood estimation. We found no evidence for a significant study block effect on AWP (type-I sum of squares: *F* = 0.29, *P* = 0.590).

To understand how tree species richness (SR) and mycorrhizal associations (MA) affected AWP in our young temperate forest plots, we fitted a structural equation model (SEM) to our data using the *piecewiseSEM* R package (*67*). Our model includes four main pathways susceptible to modulate AWP: (1) a direct effect of SR on AWP, (2) an indirect effect of SR on AWP via its effect on SSCI, (3) an indirect effect of SR on AWP via its effect on tree mortality, and (4) a direct effect of the relative proportion of AM and EM trees in the community on AWP. This piecewise SEM consisted of a set of three mixed-effects models. All models were fitted using tree species composition as a random effect. The goodness-of-fit of our SEM was assessed using Fisher’s C test statistic (*67*).

To identify the most important drivers of ΔSSCI (i.e., the net effect of tree species diversity on stand structural complexity) and its effect on wood productivity, we fitted two additional piecewise SEMs. Each SEM focused on one key functional characteristic of the species: (1) their shade tolerance (CWM_ST_, FD_ST_) or (2) their phenotypic plasticity (CWM_RDPI_, FD_RD_). For the sake of simplicity, we will refer to the community-weighted mean or the functional dispersion of a trait as CWM and FD, respectively, in what follows. In each SEM, we tested three main pathways via which SR could modulate ΔSSCI: (1) a direct effect of SR on ΔSSCI, (2) an indirect effect of SR on ΔSSCI via its effect on CWM, and (3) an indirect effect of SR on ΔSSCI via its effect on FD. The direct effect of ΔSSCI on AWP was represented in the model, as well as a direct effect of SR on AWP and a direct effect of FD on AWP. In the SEM focusing on phenotypic plasticity, we also included a direct link between CWM_RDPI_ and FD_RDPI_. Each piecewise SEM consisted of a set of two simple linear models and two mixed-effects models (tree species composition used as a random effect). The goodness-of-fit of each SEM was assessed using Fisher’s C test statistic (*67*).

All statistical analyses were performed in R 4.2.1 (*59*) using the packages *dplyr* (*68*), *ggplot2* (*69*), *ggeffects* (*70*), *lme4* (*71*), *lmerTest* (*72*), *MuMIn* (*73*), *nlme* (*74*), *variancePartition* (*75*), *VoxR* (*60*) and *sp* (*61*).

## Acknowledgments

We are grateful to Alexandra Koller and Louis Georgi for their assistance in data collection and to Dr. Martin Ehbrecht and Karl Friedrich Reich for their suggestions in data analysis. We are also thankful to Norman Döring for his technical assistance.

## Author contributions

N.E. and O.F. designed and established the experiment.

T.R., A.F., G.v.O. and H.B. conceived the idea of the study.

T.R collected the TLS data and J.Q. collected the diameter and height data.

T.R., A.F. and B.D. analysed the data and created the figures.

T.R. wrote the first draft of the manuscript with support from A.F. and B.D. All authors contributed to the discussion of the results and revisions.

---

**Table S1.** Results of linear mixed-effects models of the effects of tree species richness on stand structural complexity (SSCI) and community productivity (AWP). SD, standard deviation. The variance explained by the fixed effects alone (marginal R^2^) and by both the fixed and random effects (conditional R^2^) was calculated according to Nakagawa & Schielzeth (2013) (76).

|  | SSCI-model | AWP-model |
| --- | --- | --- |
| **Random effects** |  |  |
| SD (species composition) | 0.970 | 0.021 |
| SD (residuals) | 0.340 | 0.023 |
| **Model fit** |  |  |
| $R^2_m$ | 0.19 | 0.11 |
| $R^2_c$ | 0.91 | 0.52 |

**Table S2.** List of tree species used in the MyDiv experiment and their associated functional characteristics, including mycorrhizal associations, net primary productivity in monoculture, as well as shade tolerance and plasticity indices. Within each category of mycorrhizal association, species are ranked from most to least productive. AMF, arbuscular mycorrhizal fungi; AWP, annual wood productivity; EMF, ectomycorrhizal fungi; RDPI, relative distance plasticity index.

| **Tree species** | **Mycorrhizal association** | **AWP (m<sup>3</sup>/year) in monoculture** | **Shade tolerance index** | **RDPI** |
| --- | --- | --- | --- | --- |
| *Prunus avium* | AMF | 0.15 | 3.33 | 0.19 |
| *Acer pseudoplatanus* | AMF | 0.14 | 3.73 | 0.10 |
| *Fraxinus excelsior* | AMF | 0.10 | 2.66 | 0.20 |
| *Sorbus aucuparia* | AMF | 0.08 | 2.73 | 0.16 |
| *Aesculus hippocastanum* | AMF | 0.07 | 3.43 | 0.29 |
| *Tilia platyphyllos* | EMF | 0.16 | 4.00 | 0.31 |
| *Betula pendula* | EMF | 0.11 | 2.03 | 0.32 |
| *Carpinus betulus* | EMF | 0.10 | 3.97 | 0.34 |
| *Quercus petraea* | EMF | 0.04 | 2.73 | 0.53 |
| *Fagus sylvatica* | EMF | 0.04 | 4.56 | 0.54 |

**Fig. S1.**
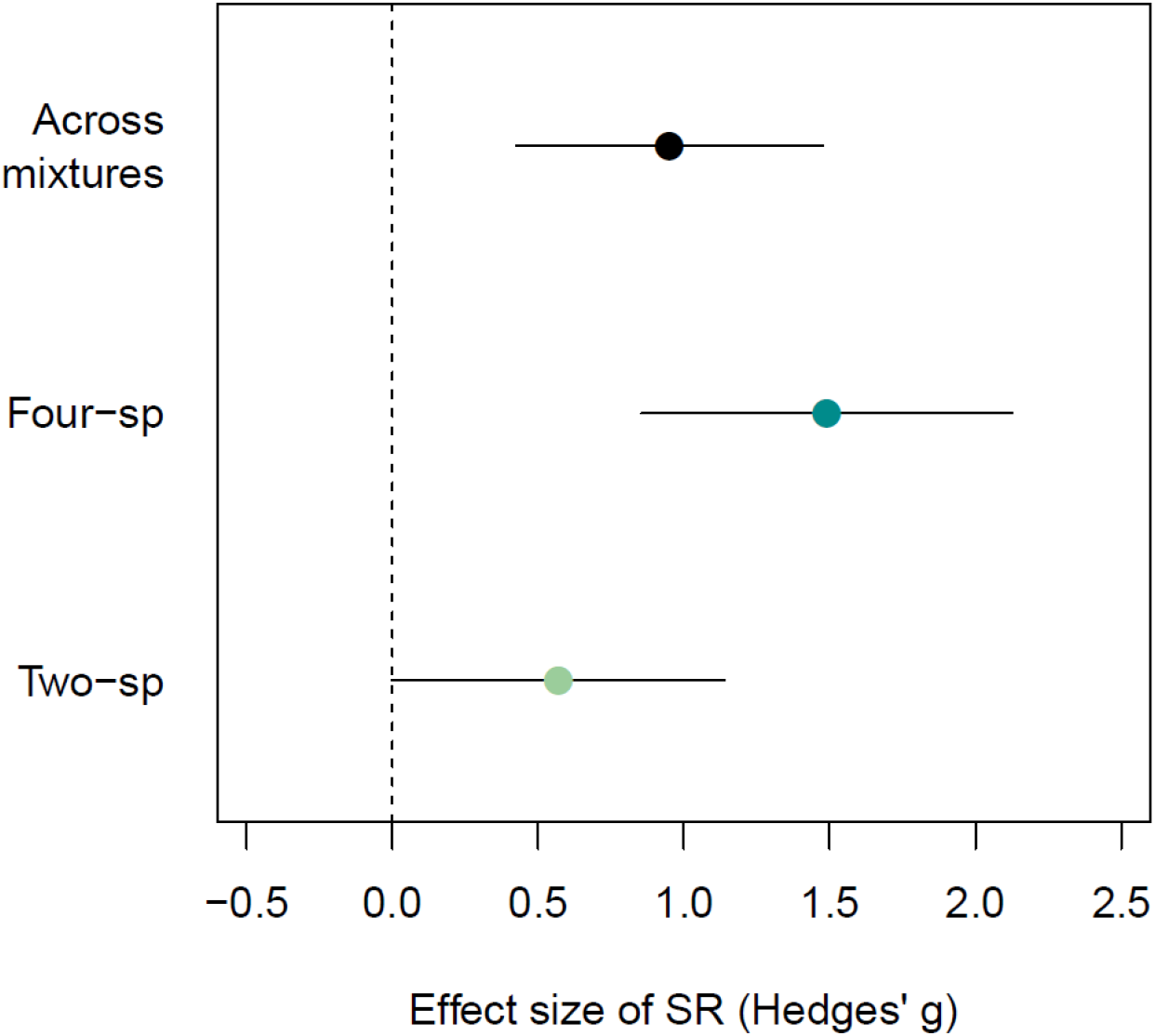
Effect sizes for the biodiversity-complexity relationship showing how tree species richness affects the strength of biodiversity effects (Hedges’ g effect size) on stand structural complexity (SSCI). Dots are predicted means of mixed-effects models, and error bars denote the 95% confidence intervals. Positive values indicate a higher structural complexity in mixed-species communities compared to monocultures, while negative values indicate the opposite. Error bars not overlapping with zero indicate significant biodiversity effects, and vice versa. Across mixtures: 2-and 4-species mixtures.

**Fig. S2.**
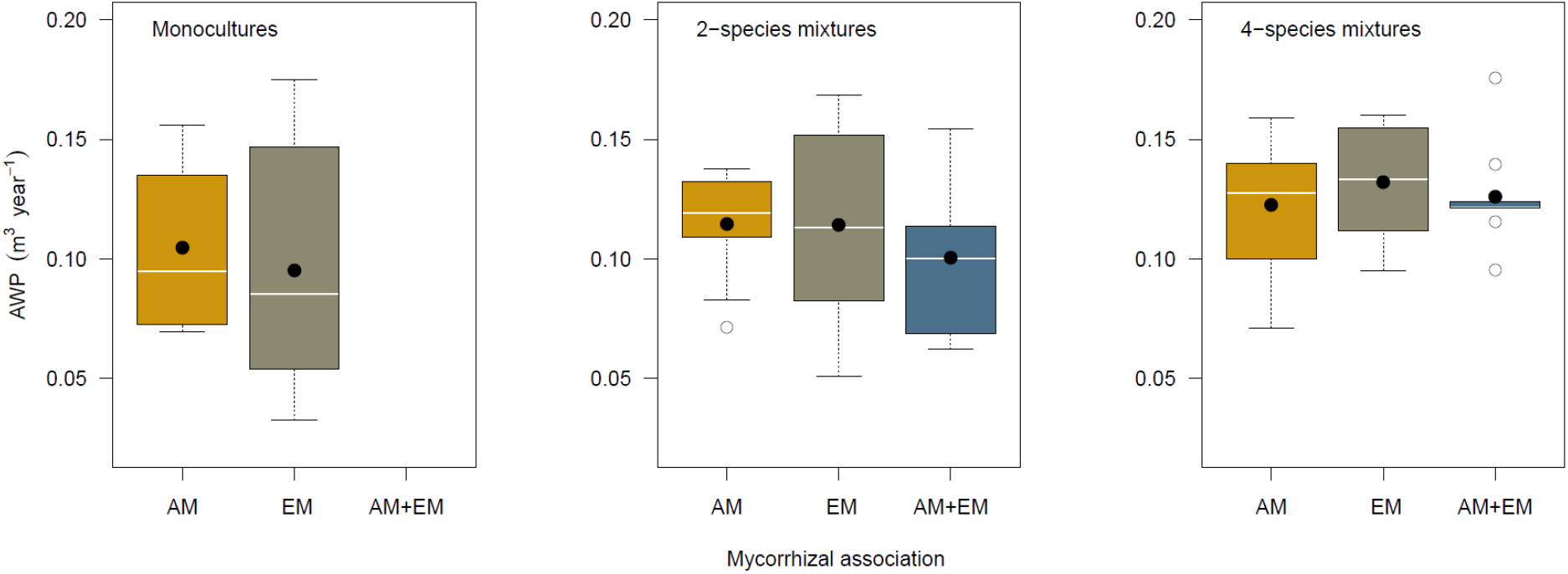
Effects of mycorrhizal associations on community productivity does not depend on tree species richness. Boxplots show the median (horizontal white lines), the mean (black dots), the 25% and 75% percentiles (edges of the box) and 1.5 times the interquartile range (whiskers) of observed community productivity. Open circles indicate productivity values that are greater or smaller than 1.5 times the interquartile range. Differences among mycorrhizal associations were not statistically significant (Tukey-Test: *P* > 0.10). Note that the interaction between tree species richness and mycorrhizal associations was not significant (*P* = 0.56). AM: arbuscular mycorrhizal tree species; EM: ectomycorrhizal tree species.

**Fig. S3.**
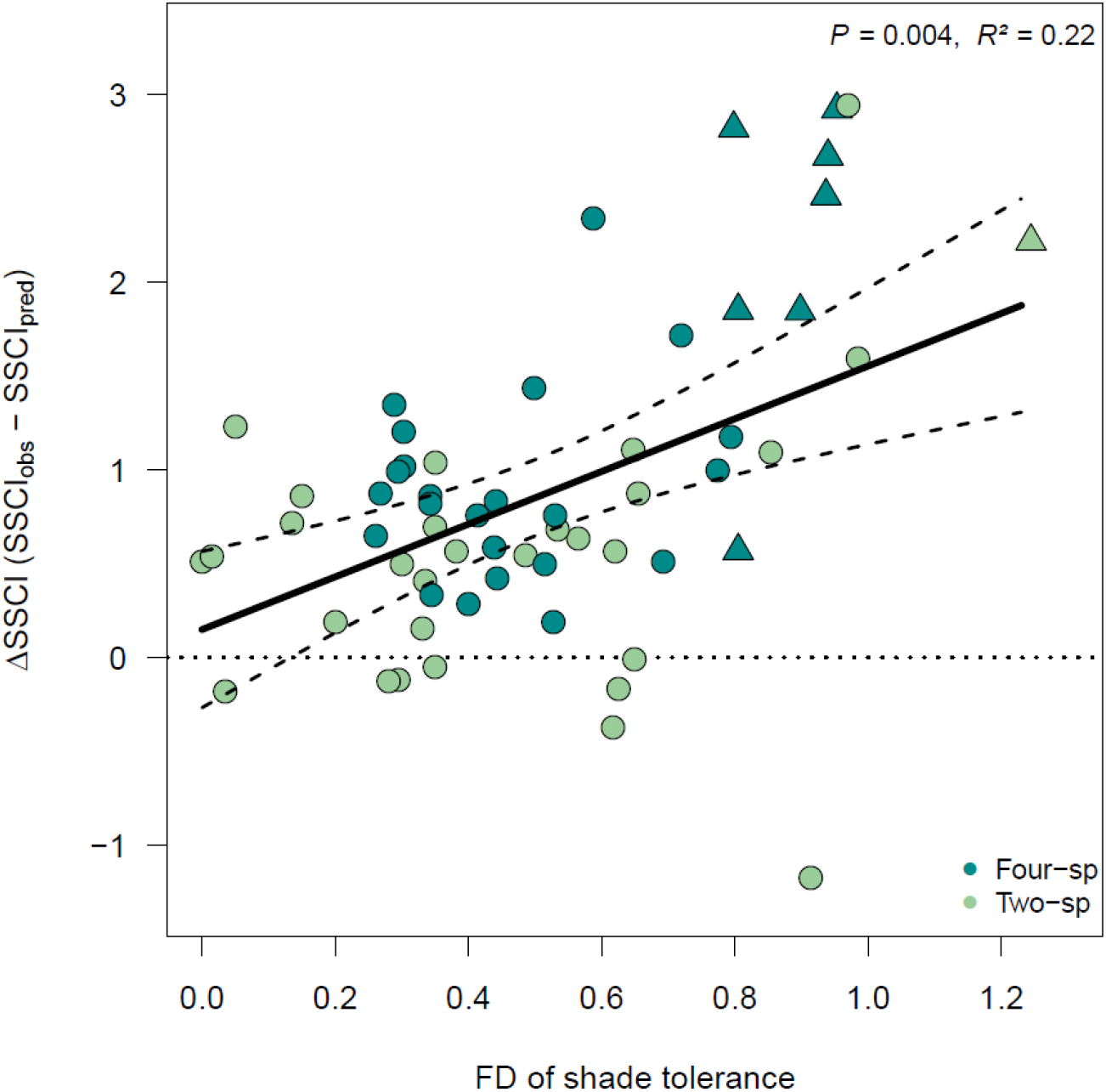
Shade tolerance-structural complexity relationship. Relationship between the net effect of tree species richness on structural complexity (ΔSSCI) and the functional dispersion (FD) of shade tolerance within mixtures. For each mixture, ΔSSCI was calculated as the difference between observed (SSCI_obs_) and predicted (SSCI_pred_) structural complexity (see Methods). The solid line is a mixed-effect model fit. The dotted lines indicate the 95% confidence interval of the prediction. Raw data are shown with individual data points, with triangles showing plots containing both *Betula pendula* and *Fagus sylvatica* growing in 2-species (light green) or 4-species (dark green) mixtures. The *R*²-value refers to the proportion of variance explained by FD.

**Fig. S4.**
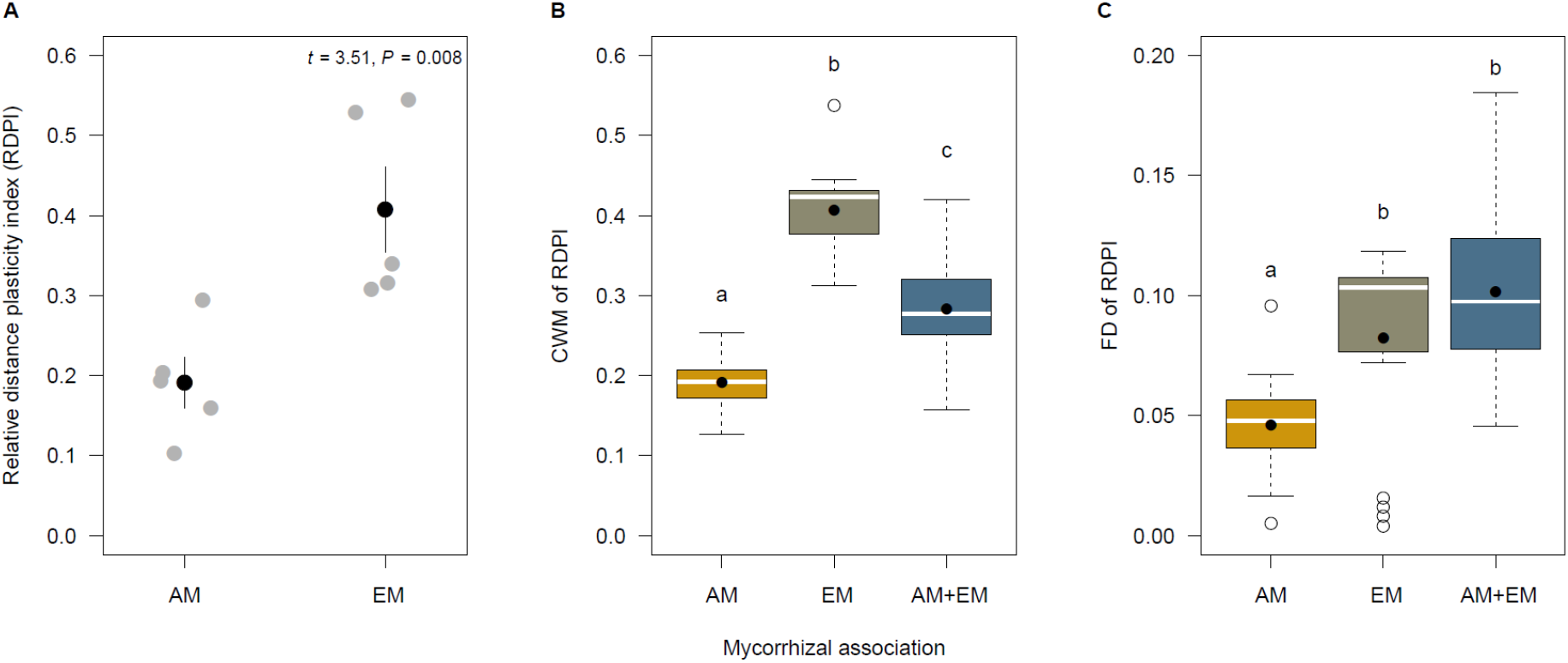
Effects of mycorrhizal association on phenotypic plasticity. Species-specific variation in mean (± standard error) relative distance plasticity index (RDPI) between arbuscular mycorrhizal (AM) and ectomycorrhizal (EM) tree species. Grey dots correspond to the RDPI values of our ten study species and error bars indicate the standard error **(A)**. Changes in community-weighted means (CWM; **B**) and functional dispersion (FD; **C**) of phenotypic plasticity of our 60 tree communities with mycorrhizal association (AM, EM, AM+EM). Boxplots show the median (horizontal black lines), the mean (black dots), the 25% and 75% percentiles (edges of the box) and the variation (error bars: 1.5× the interquartile range) of observed community productivity. Open circles indicate values that are greater or smaller than 1.5 times the interquartile range. Different letters indicate significant differences among mycorrhizal types (Tukey-Test: *P* ≤ 0.5).

**Fig. S5.**
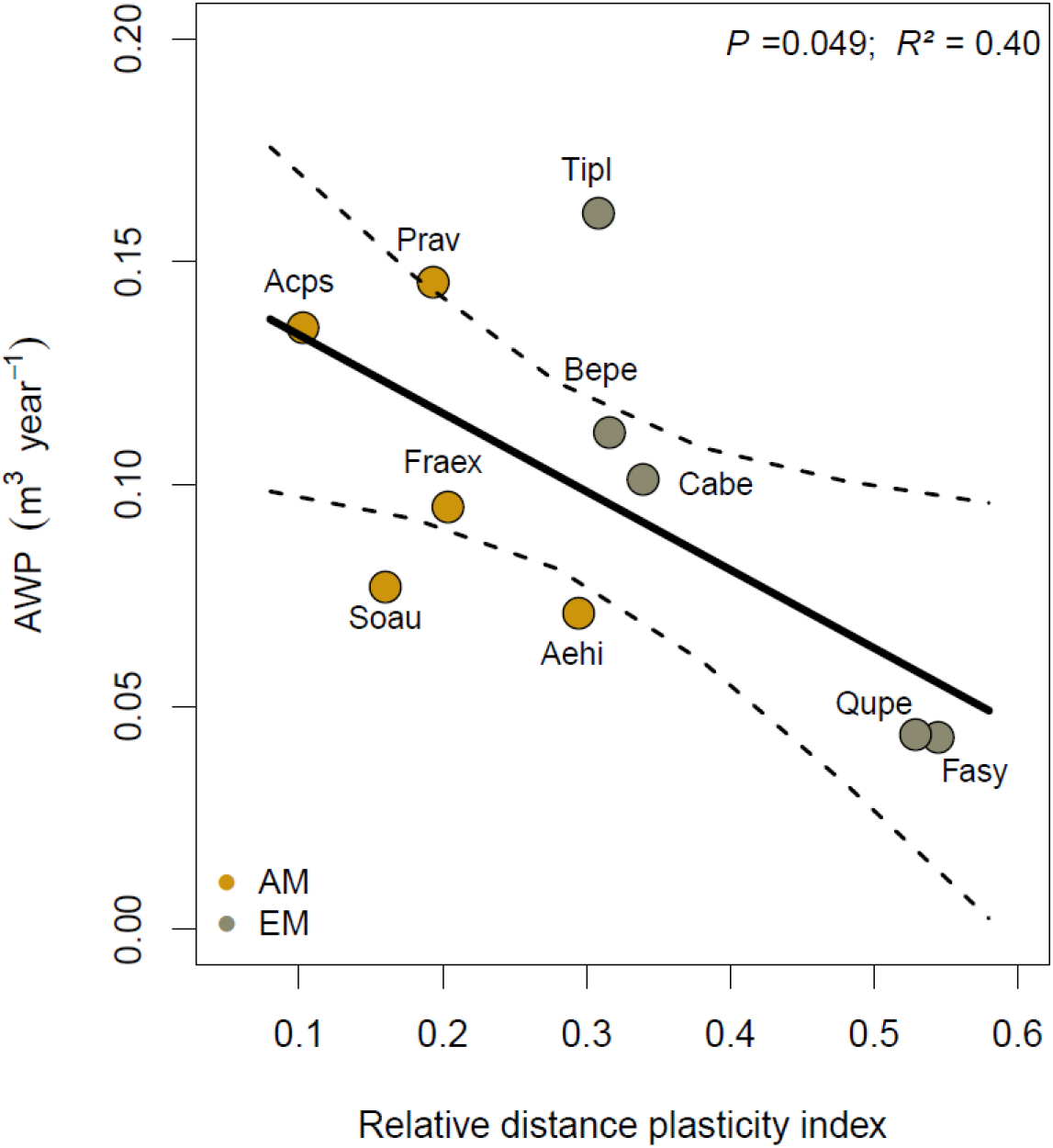
Relationship between phenotypic plasticity and wood productivity (AWP) at the species level. The solid line is a linear model fit and the dotted lines indicate the 95% confidence interval. Dots represent the observed relative distance plasticity indices (RDPI) of the ten study species. The *R*²-value indicates the proportion of variance explained by species-specific RDPI. AM: arbuscular mycorrhizal tree species; EM: ectomycorrhizal tree species

**Fig. S6.**
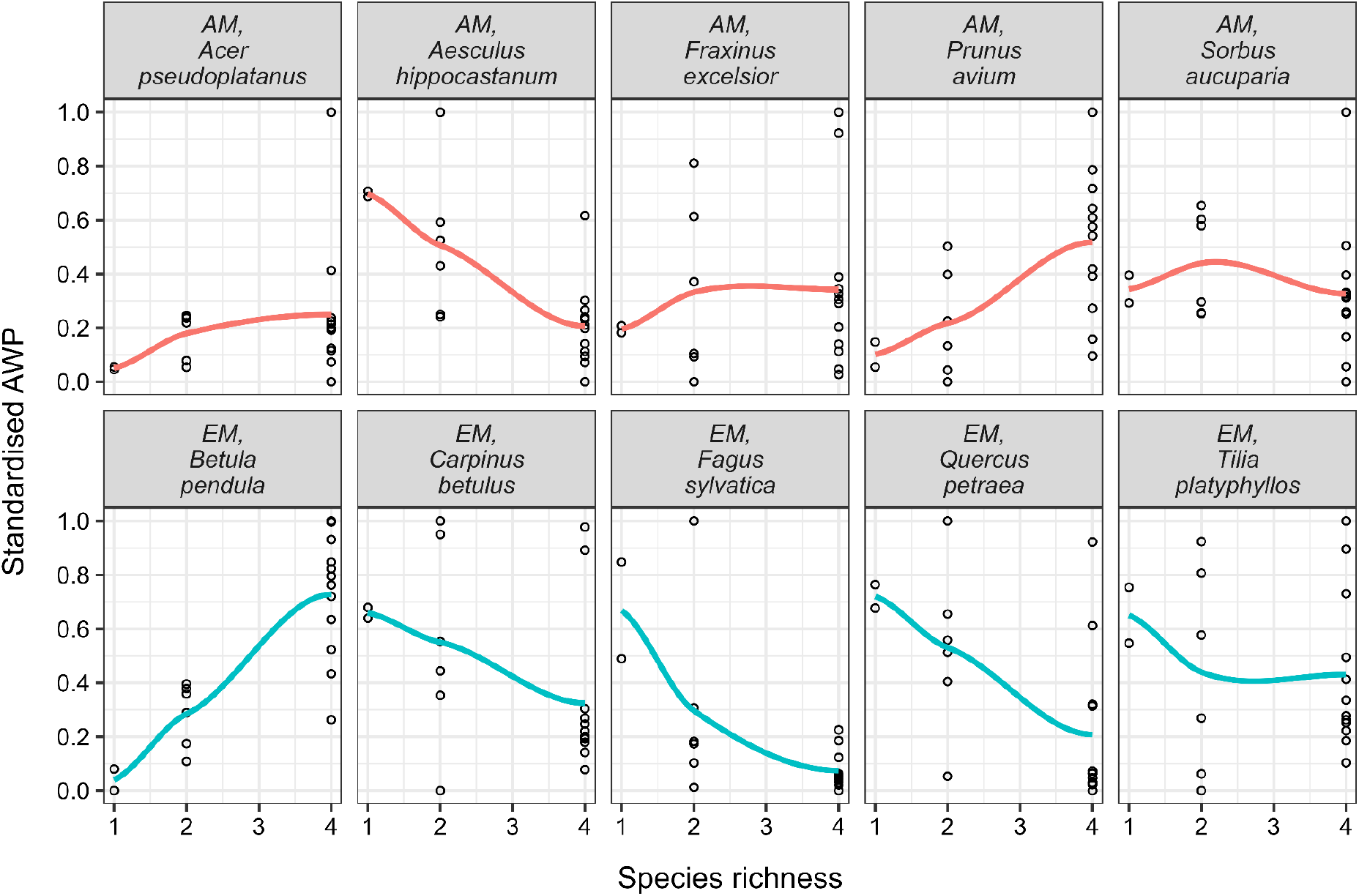
Species-specific effects of tree species richness on wood productivity (AWP). For each species, wood productivity data were rescaled using a min-max normalisation procedure. In our paper, we assessed tree phenotypic plasticity by quantifying the extent to which AWP of each species changes along the species richness gradient. We expect tree species with low phenotypic plasticity to show small differences in AWP between monoculture and mixtures of two and four species. Tree species with high phenotypic plasticity, however, would show large differences in AWP across species richness levels. In each panel, lines are loess smooths. Species are grouped per mycorrhizal association (AM, arbuscular mycorrhizal trees; EM, ectomycorrhizal trees).

**Fig. S7.**
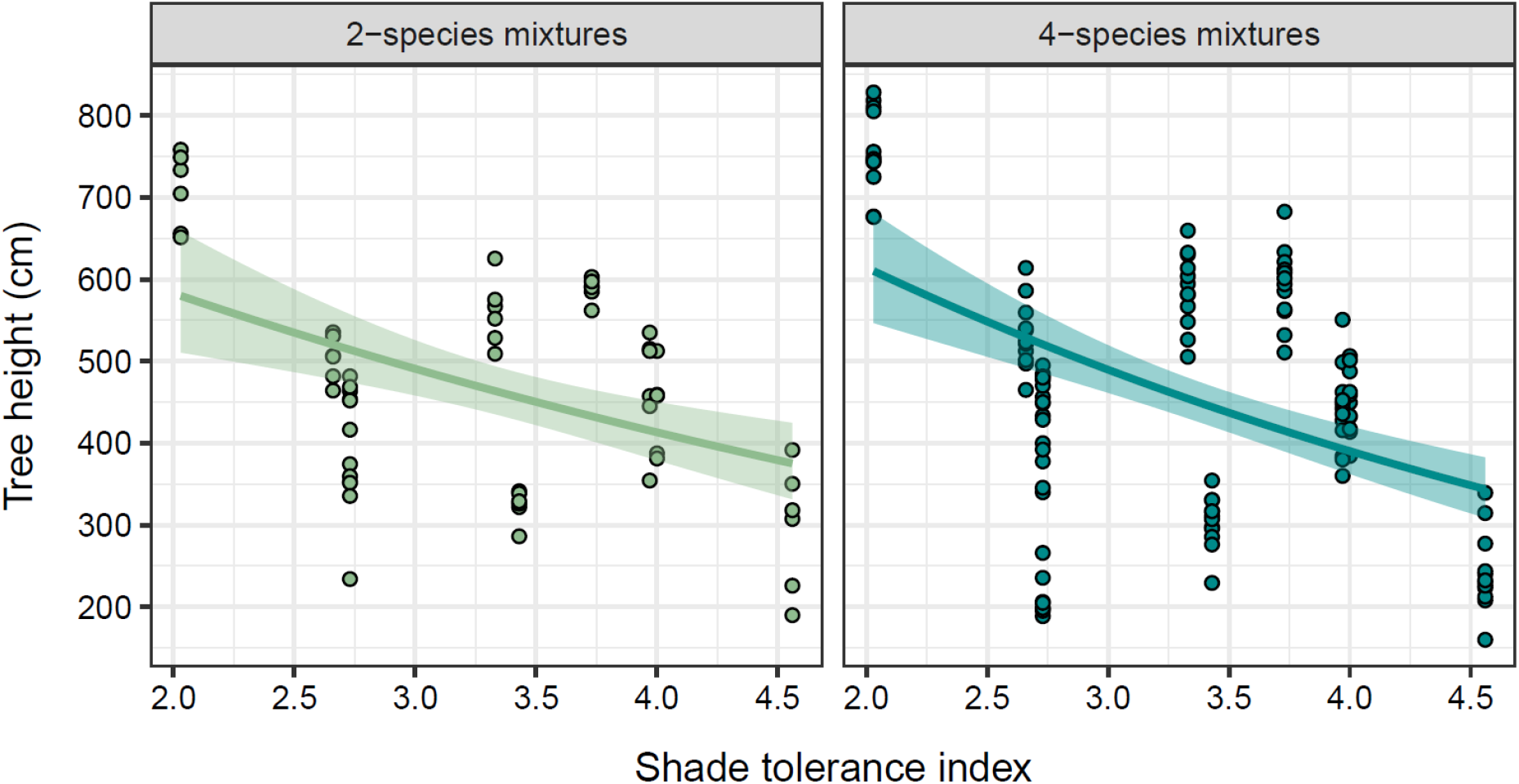
Effects of shade tolerance on tree height growth. Univariate relationships are displayed for 2-species mixtures and 4-species mixtures. Individual dots represent the average height in 2021 (year of scanning) of a species in a plot. Lines are generalised linear model fits (Gamma distribution and log link function). Shade tolerance was used as a continuous variable in the model, while species richness was used as a factor with two levels. There was no interaction between tree species richness and shade tolerance (P = 0.377), but shade tolerance had a strong negative effect on tree height (P = 0.0008). Across all species included in our experiment, tree species richness did not affect tree height (P = 0.441).

**Fig. S8.**
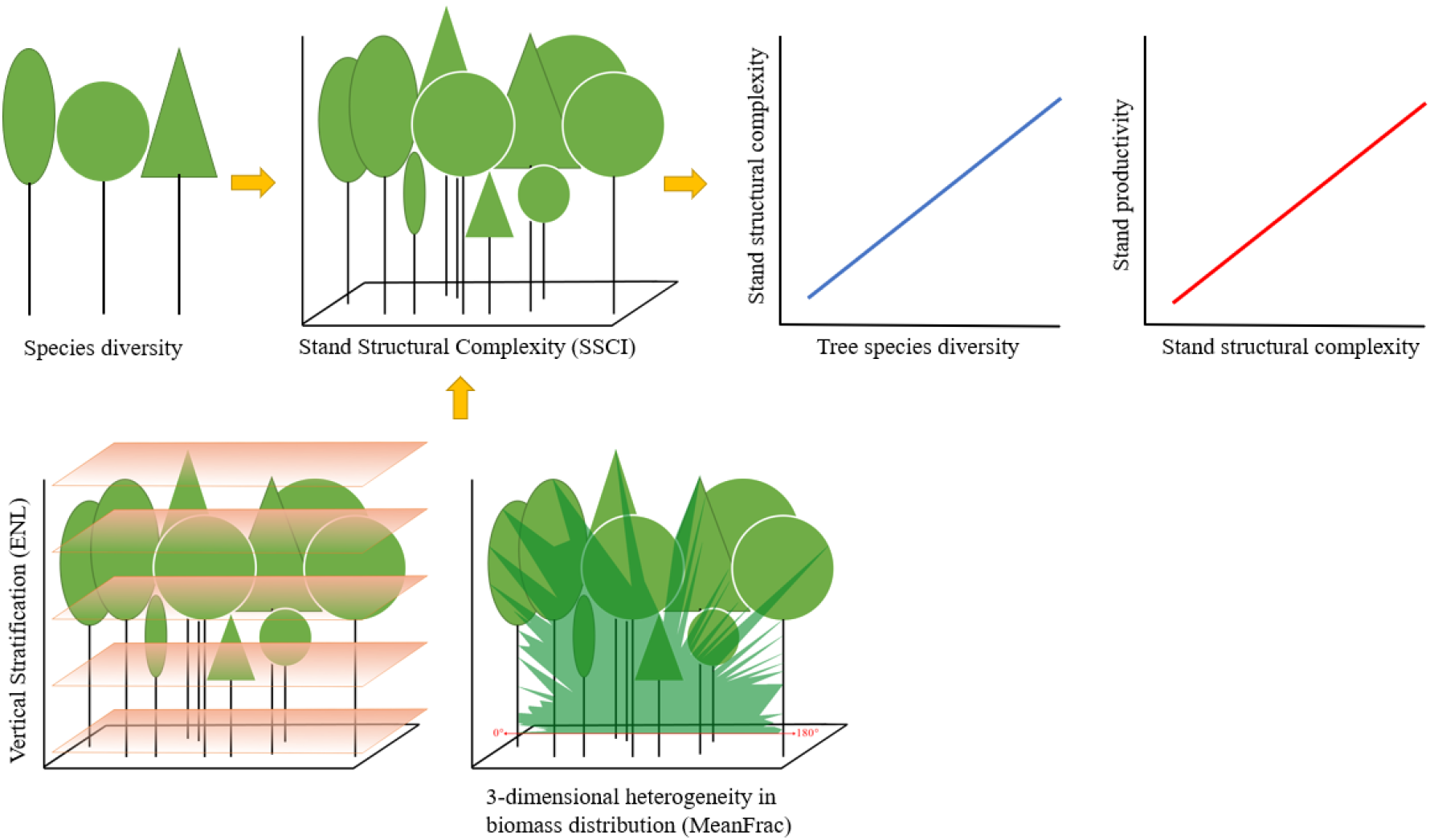
Graphical representation of tree species diversity effects on stand productivity, enhanced by stand structural complexity. Greater tree species richness allows different species to grow in different strata depending on their degree of morphological, architectural and functional dissimilarity. This promotes greater vertical stratification (ENL) and 3-dimensional heterogeneity in biomass distribution (MeanFrac) of the stand, two basic components of stand structural complexity. Therefore, we hypothesised that the structural complexity of a tree community enhances resource acquisition of the trees and helps to store wood biomass in different strata of the stand.

